# Microtubule-independent movement of the fission yeast nucleus

**DOI:** 10.1101/2020.08.29.273409

**Authors:** Sanju Ashraf, David A. Kelly, Kenneth E. Sawin

**Affiliations:** Wellcome Centre for Cell Biology, School of Biological Sciences, University of Edinburgh, Michael Swann Building, Max Born Crescent, Edinburgh EH9 3BF, UK

## Abstract

Movement of the cell nucleus typically involves the cytoskeleton and either polymerization-based pushing forces or motor-based pulling forces. In fission yeast *Schizosaccharomyces pombe*, nuclear movement and positioning are thought to depend on microtubule polymerization-based pushing forces. Here we describe a novel, microtubule-independent, form of nuclear movement in fission yeast. Microtubule-independent nuclear movement is directed towards growing cell tips, and it is strongest when the nucleus is close to a growing cell tip, and weakest when the nucleus is far from that tip. Microtubule-independent nuclear movement requires actin cables but does not depend on actin polymerization-based pushing or myosin V-based pulling forces. Vesicle-associated membrane protein (VAMP)-associated proteins (VAPs) Scs2 and Scs22, which are critical for endoplasmic reticulum-plasma membrane contact sites in fission yeast, are also required for microtubule-independent nuclear movement. We also find that in cells in which microtubule-based pushing forces are present, disruption of actin cables leads to increased fluctuations in interphase nuclear positioning and subsequent altered septation. Our results suggest two non-exclusive mechanisms for microtubule-independent nuclear movement, which may help illuminate aspects of nuclear positioning in other cells.

## INTRODUCTION

Movement and positioning of the cell nucleus depends on cell type, cell-cycle stage, and cell migration and differentiation states (Bone and Starr, 2016; Dupin and Etienne-Manneville, 2011; Gundersen and Worman, 2013; Lele et al., 2018; Xiang, 2018). Precise nuclear positioning is important in cell division, cell migration, and development, and in humans, mispositioning of the nucleus is associated with several different pathologies, including muscular dystrophy and cardiomyopathy, lissencephaly and premature ageing (Gundersen and Worman, 2013; Isermann and Lammerding, 2013). In budding yeast *Saccharomyces cerevisiae*, nuclear positioning is required for the daughter cell to inherit a nucleus from the mother cell (Moore and Cooper, 2010; Pruyne et al., 2004), and in the cylindrically-shaped fission yeast *Schizosaccharomyces pombe*, nuclear positioning is required to generate equal-sized daughter cells during cell division (Chang and Nurse, 1996; Daga and Chang, 2005; Tolic-Norrelykke et al., 2005).

In organisms from humans to yeasts, nuclear movement generally involves proteins of the cytoskeleton and protein complexes associated with the nuclear envelope (NE) (Gundersen and Worman, 2013; Tapley and Starr, 2013). Both microtubules (MTs) and actin filaments have been implicated in generating forces for movement (Dupin and Etienne-Manneville, 2011; Reinsch and Gonczy, 1998); in some instances, intermediate filaments may also contribute (Dupin et al., 2011). In addition, in many cells, LINC (linkers of nucleoskeleton and cytoskeleton) protein complexes, which span the inner and outer nuclear membranes, help transmit forces from the cytoskeleton to the nucleus (Burke, 2019; Crisp et al., 2006; Padmakumar et al., 2005; Tapley and Starr, 2013).

MT-dependent nuclear movement can occur via multiple mechanisms. In some cases, the nucleus is closely associated with a microtubule organizing center (MTOC; e.g. the centrosome in animal cells, or the spindle pole body (SPB) in yeast), and movement of both the MTOC and the associated nucleus depends on dynamic changes in the MT network, driven variously by microtubule polymerization, depolymerization, and/or cortex-localized MT motors (e.g. dynein). Examples of this include movement of the male pronucleus in *Xenopus* early development (Reinsch and Gonczy, 1998), the *Drosophila* oocyte nucleus during dorsal/ventral axis specification (Zhao et al., 2012), and fibroblast nuclei during cell migration (Levy and Holzbaur, 2008). In other cases, the nucleus itself moves along MTs, driven by motor proteins (e.g. kinesins or dyneins) associated with the NE. Examples include movement of the female pronucleus during *Xenopus* development (Reinsch and Gonczy, 1998), a variety of nuclear movements in *Caenorhabditis elegans* (Fridolfsson and Starr, 2010), and interkinetic nuclear migration of neuronal precursors during vertebrate brain development (Bertipaglia et al., 2018; Tsai et al., 2010).

Actin-dependent nuclear movement generally depends on forces that push the nucleus, as a result of actin polymerization, actin network flow and/or actomyosin cytoskeleton/contractility. In some cases, these work in concert with microtubule-dependent forces. Actin-dependent nuclear movements are seen in migrating fibroblasts and astrocytes (Dupin et al., 2011; Gomes et al., 2005; Luxton et al., 2010), in certain types of neuroepithelia (Yanakieva et al., 2019), and in nurse cells in the *Drosophila* ovary (Huelsmann et al., 2013). Currently there is little evidence for NE-associated motor proteins driving nuclear movement along actin filaments, at least in animal cells. However, in *Arabidopsis thaliana*, nuclear movement involves a plant-specific myosin, myosin XI-i (also known as Myo11G), which localizes to the NE and links the nucleus to actin filaments (Tamura et al., 2013).

In budding yeast, movement of the nucleus from the mother cell into the daughter bud neck involves MTs directly and actin filaments indirectly (Moore and Cooper, 2010; Pruyne et al., 2004). Initially, MTs emanating from one of the duplicated SPBs on the NE are guided along actin filaments into the bud neck, via association of MT plus-ends with the class V myosin Myo2. Subsequently, a combination of MT depolymerization at cortical MT-contact sites in the daughter cell and cortical dynein-driven MT sliding provides force to pull the nucleus into the bud neck.

In fission yeast, the interphase nucleus is continuously maintained at the cell center as the cell grows by elongation at its tips. The position of the late-interphase/early-mitotic nucleus defines the future plane of cell division, and nuclear mispositioning leads to unequal segregation of cellular components and unequal-sized daughter cells (Chang and Nurse, 1996; Daga and Chang, 2005; Tolic-Norrelykke et al., 2005). Because interphase growth in fission yeast is normally asymmetric (i.e. monopolar early in the cell cycle, and bipolar later in the cell cycle; (Mitchison and Nurse, 1985)), the precision of nuclear centering has suggested that an active mechanism positions the nucleus, and experiments using tubulin mutants, MT-depolymerizing drugs, cell centrifugation and optical tweezers have indicated that MTs are critical for nuclear centering (Daga et al., 2006; Hagan and Yanagida, 1997; Sawin et al., 2004; Toda et al., 1983; Tolic-Norrelykke et al., 2005; Tran et al., 2001; Umesono et al., 1983). During interphase, fission yeast MTs exist as several independent antiparallel bundles that undergo repeated cycles of polymerization and depolymerization (Sawin and Tran, 2006), and these MTs can exert pushing forces on the nucleus during MT polymerization, leading to a model in which MT pushing forces drive nuclear positioning (Tran et al., 2001). Consistent with this model, the fission yeast SPB, which is associated with the nucleus, exhibits MT-dependent oscillations parallel to the long axis of the cell, and the NE similarly shows MT-dependent deformations (Tran et al., 2001).

Although the idea that MT-dependent forces position the fission yeast nucleus has been accepted for many years, the question of whether the nucleus can still move in a cell that lacks MTs has, surprisingly, never been investigated directly. In this work, using live-cell imaging and a combination of drug perturbations and mutants, we show that robust movement of the fission yeast nucleus persists even after MT depolymerization. This MT-independent nuclear movement is directed towards growing cell tips, and we further show that it depends on actin cables nucleated from growing tips, but not on myosin motors. We also find that MT-independent nuclear movement requires proper spatial organization of the endoplasmic reticulum (ER), which in yeast can link the NE with the plasma membrane. Overall, MT-independent movement of the fission yeast nucleus appears to involve mechanisms of force generation that are distinct from those previously described for nuclear movement. Finally, we show that in conjunction with MT-dependent pushing forces, the mechanisms underlying MT-independent nuclear movement contribute to accurate positioning of the nucleus at the cell center.

## RESULTS

### Unidirectional nuclear movement after microtubule depolymerization

To investigate whether movement of the fission yeast nucleus can occur in the absence of MTs, we depolymerized MTs and imaged cells during interphase growth (cell elongation). We used a Cdc42/Rac Interactive Binding (CRIB)-3mCitrine fusion protein (Mutavchiev et al., 2016), which binds to the active (GTP-bound) form of Cdc42, as a reporter of both cell growth and nuclear position (**Fig. 1A**). CRIB-3mCitrine localizes to growing cell tips during interphase and to the septum during cell division, and it is also present in the nucleus throughout the cell cycle, although nuclear localization is artifactual and unrelated to Cdc42 function (Bendezu et al., 2015; Mutavchiev et al., 2016; Tatebe et al., 2005). We depolymerized MTs with the drug methyl benzimidazol-2-yl carbamate (MBC). Consistent with previous work (Sawin and Snaith, 2004), in cells expressing GFP-tubulin, nearly all MTs disappeared within 15 minutes after MBC addition (**Fig. 1B**). The only remaining MTs were short “stubs” that would be unable to mediate pushing forces for MT-dependent nuclear centering, and even these stubs were barely visible after 20-30 minutes. We analyzed nuclear movement specifically in monopolar-growing cells, i.e. cells growing at only one tip. As shown in **Fig. 1C**, this is because in bipolar-growing cells, the nucleus would be expected to be in the cell center not only when there is an active mechanism positioning the nucleus to the cell center but also when there is no nuclear movement at all. By contrast, in monopolar-growing cells, an active nuclear positioning mechanism would result in unidirectional movement of the nucleus over time, in the same direction as growth of the growing cell tip.

**Figure 1.**
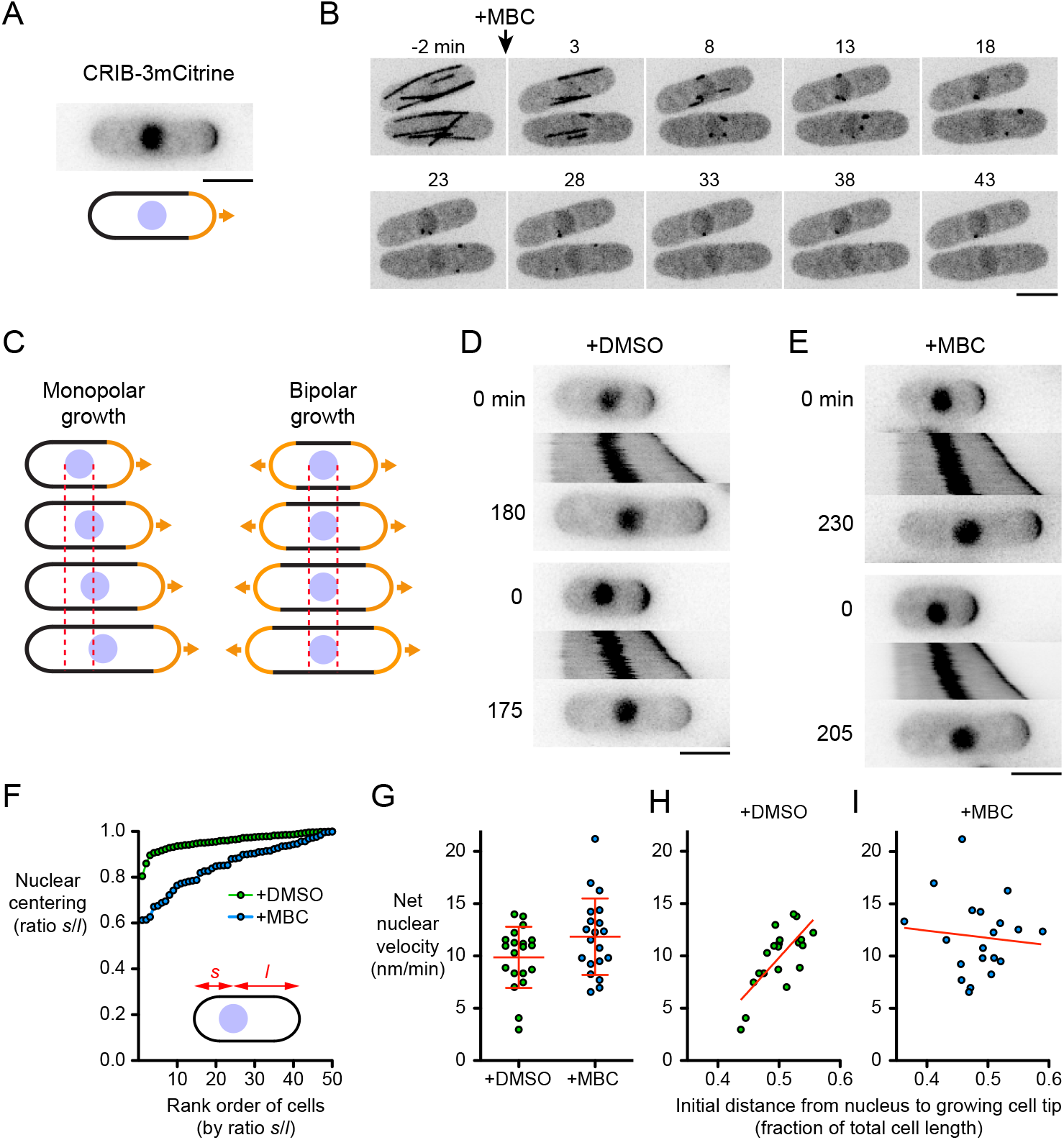
Microtubules are not required for nuclear movement in fission yeast. **(A)** CRIB-3mCitrine as a reporter of growing cell tips and nuclear position. **(B)** Timepoints from movie showing kinetics of interphase microtubule (GFP-Atb2) depolymerization after treatment with 25 µg/mL methyl benzimidazol-2-yl carbamate (MBC). **(C)** Cartoon showing how nuclear movement is most easily assayed in monopolar-growing cells rather than bipolar-growing cells. Dashed lines indicate boundaries of nucleus at time zero. See text for details. **(D**,**E)** Timepoints and kymographs of nuclear movement from movies of DMSO-treated (D) and MBC-treated (E) cells. Timepoints correspond to beginning and end of kymographs (min). Two examples are shown for each condition. **(F)** Nuclear centering in DMSO-treated and MBC-treated cells. Cells are ordered by ratio *s*/*l*, in which *s* and *l* are the shorter and longer distances (respectively) from nucleus to cell tip (50 cells measured for each condition; see Materials and Methods). **(G)** Net nuclear velocity towards growing cell tips in monopolar-growing DMSO-and MBC-treated cells. Red lines show mean and SD (n=20 for each condition). Difference was not statistically significant (*t*-test, *p*=0.067) **(H**,**I)** Net nuclear velocity in DMSO-treated (H) and MBC-treated (I) cells, plotted against distance from nucleus to growing cell tip (as a fraction of total cell length) at beginning of movie. Data are from the same cells as in G. Red lines show linear regression (in H, *r*2=0.53, *p*=0.0003; in I, *r*2=0.009, *p*=0.69). Bars, 5 µm.

In control (DMSO-treated) monopolar-growing cells, the nucleus moved in the same direction as the growing cell tip in 100% of cells (100 out of 100 cells**; Fig. 1D, Movie 1**). Consistent with previous work (see Introduction), in these cells the nucleus was almost always near the cell center **(Fig. 1 F)**. Interestingly, in MBC-treated monopolar-growing cells, the nucleus also moved in the same direction as the growing cell tip, in 100% of cells (150 out of 150 cells) (**Fig. 1E, Movie 1)**. We will refer to this movement as “MT-independent nuclear movement” (MINM).

The net velocity of MINM (∼12 nm/min) was only slightly different from nuclear movement in control cells (i.e. not statistically significant; **Fig. 1G**). This was surprising, given the strong evidence implicating MTs in fission yeast nuclear positioning (see Introduction). However, fidelity of nuclear centering in MBC-treated cells was significantly decreased compared to control cells (**Fig. 1F**). Although similarities in nuclear velocity would seem to be incompatible with differences in nuclear centering, we reconciled these results by analyzing the relationship between nuclear velocity and the position of the nucleus at the start of imaging (**Fig. 1H, I**). In control cells, net nuclear velocity towards the growing cell tip was lowest when the nucleus was close to the growing cell tip (i.e. at the start of imaging) and highest when the nucleus was far from the growing tip (r^2^=0.53; p=0.0003). This is consistent with MTs providing a homeostatic nuclear-centering mechanism (Tran et al., 2001). By contrast, in MBC-treated cells, there was no specific relationship between nuclear velocity and the position of the nucleus at the start of imaging (r^2^=0.01; p=0.69),suggesting that MINM is largely independent of a centering mechanism. Collectively, these results indicate that nuclear movement in fission yeast does not depend exclusively on force generation by MTs; MT-independent mechanisms are also present. However, highly accurate centering of the nucleus requires MTs.

### Microtubule-independent nuclear movement (MINM) requires actin cables

We hypothesized that MINM may depend on the actin cytoskeleton. During fission yeast interphase, actin filaments are present in two distinct structures: cortical actin patches and actin cables (Kovar et al., 2011; Mishra et al., 2014)(**Fig. S1**). Cortical actin patches are nucleated by the Arp2/3 complex, primarily at cell tips, and are involved in endocytosis. Actin cables, which contain multiple actin filaments, are nucleated by formin For3 at growing cell tips and are involved in intracellular transport. Because cortical actin patches are far away from the cell nucleus, they would not be expected to directly participate in MINM. By contrast, actin cables extend from growing cell tips into the cell interior and can easily contact the nucleus. We therefore investigated a role for actin cables in MINM.

To test whether MINM requires actin cables, we used *for3Δ* deletion mutants (see Materials and Methods). In *for3Δ* cells, interphase actin cables have been described as being either severely compromised or absent altogether, although polarized growth still occurs (Feierbach and Chang, 2001; Nakano et al., 2002). Using Lifeact-mCherry (Huang et al., 2012; Riedl et al., 2008), we observed a small number of residual actin cables in *for3Δ* cells, but these were shorter, fainter, and fewer in number than in wild-type cells (**Fig. S1**). Residual actin cables may be due to low-level activity of formin Cdc12, normally involved in cytokinesis (Burke et al., 2014). We analyzed monopolar-growing *for3Δ* cells as described above for wild-type cells (**Fig. 2A**,**B**,**G; Movie 2**). In DMSO-treated *for3Δ* cells, the nucleus moved in the same direction as the growing cell tip in 100% of cells (8 out of 8 cells; the number of cells was low because most untreated and DMSO-treated *for3Δ* cells showed premature bipolar growth; see Materials and Methods), By contrast, in MBC-treated *for3Δ* cells, the nucleus remained stationary in 91% of cells (75 out of 82 cells); the remaining 9% of cells showed slight movement, but this did not resemble MINM. These results indicate that For3 is required for MINM. Although *for3Δ* cells may contain a small number of actin cables, these are insufficient and/or unable to support MINM.

**Figure 2.**
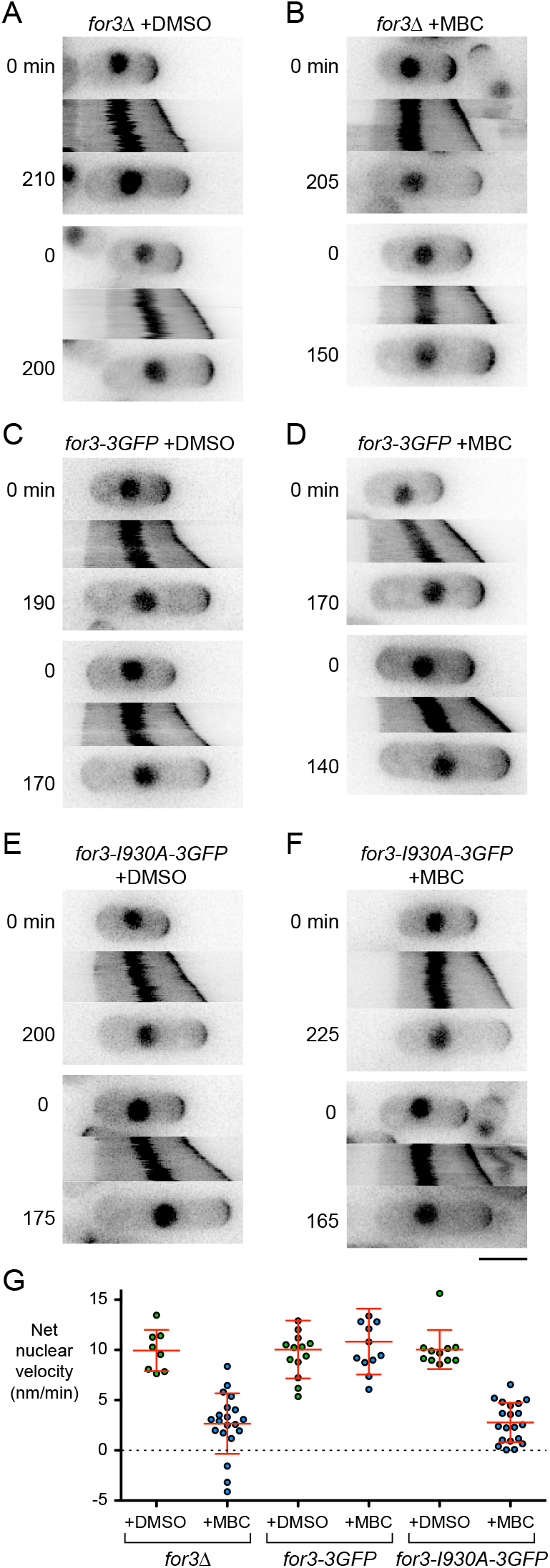
Microtubule-independent nuclear movement (MINM) requires actin cables. **(A-F)** Timepoints and kymographs of nuclear movement from movies of cells with the indicated genotypes and treatments. Timepoints correspond to beginning and end of kymographs (min). Two examples are shown for each genotype and condition. **(G)** Net nuclear velocity towards growing cell tips in monopolar-growing cells for the genotypes and treatments indicated. Red lines show mean and SD. Non-zero mean velocities for *for3Δ*+MBC and for *for3-I930A-3GFP*+MBC are likely an artifact of gradual swelling at cell non-growing tips, combined with the method used to determine net nuclear velocity (see Materials and Methods). Bar for A-F, 5 µm.

We next tested whether For3’s role in MINM is specifically related to its role in actin nucleation. For3 contains several domains, with distinct functions and binding partners (Goode and Eck, 2007; Kovar, 2006; Martin and Chang, 2006; Martin et al., 2005; Martin et al., 2007). We analysed *for3-I930A* mutants, which contain a single point mutation in the actin-binding region of the Formin Homology 2 (FH2) domain, directly involved in actin nucleation (Martin and Chang, 2006; Xu et al., 2004). We assayed nuclear movement in monopolar-growing *for3-I930A-3GFP* cells and *for3-3GFP* control cells after DMSO and MBC treatment (**Fig. 2C-G**; **Movie 3**; note that although both For3-3GFP and For3-I930A-3GFP localize to cell tips (Martin and Chang, 2006), they are too faint to be seen relative to CRIB-3mCitrine). In DMSO-and MBC-treated *for3-3GFP* cells, the nucleus moved in the direction of the growing cell tip in 100% of cells (50 out of 50 and 60 out of 60 cells, respectively). DMSO-treated *for3-I930A-3GFP* cells showed similar behavior (11 out of 11 cells; the number of cells was low because, like *for3Δ* cells, *for3-I930A-3GFP* cells normally showed premature bipolar growth; see Materials and Methods). By contrast, in MBC-treated *for3-I930A-3GFP* cells the nucleus was stationary in 91% of cells (78 out of 86 cells); as with *for3Δ* cells, the remaining 9% of cells showed slight movement, but this did not resemble MINM. We conclude that For3-mediated actin cable nucleation is critical for MINM.

### MINM depends on proximity of the nucleus to the growing cell tip

Because MINM is directed towards the growing cell tip and is dependent on actin cables, which are nucleated by For3 at the growing cell tip, we hypothesized that MINM may involve a mechanical connection between the nucleus and actin cables extending from the growing tip. If so, then MINM might be expected to be strongest when the nucleus is closest to the growing tip (i.e. where the local concentration of actin cables should be highest), and weakest when the nucleus is far away from the growing tip.

We tested this by treating cells with MBC and then centrifuging them to displace the nucleus randomly towards either cell tip, followed by imaging in the presence of MBC (**Fig. 3A**). When cells are centrifuged in the absence of MBC, the nucleus returns to the cell center within 20 minutes after centrifugation (Daga et al., 2006). Interestingly, we found that in the presence of MBC, the nucleus behaved very differently, and the type of behavior observed depended on whether centrifugation displaced the nucleus towards the growing or the non-growing cell tip. When displaced towards the growing cell tip, the nucleus continued to move towards the growing tip (i.e., away from the cell center), and nuclear velocity was greatest when centrifugation displaced the nucleus very close to the growing tip (**Fig. 3B**,**D; Movie 4**). However, when displaced towards the non-growing tip, the nucleus showed essentially no movement (**Fig**.**3C**,**D**; **Movie 4**). Intermediate displacements led to a range of nuclear velocities, all towards the growing tip. These results are consistent with a model in which MINM is most efficient when the nucleus is in a region of the cell containing a high local concentration of actin cables (see Discussion). Conversely, when the nucleus is “out of reach” of actin cables, MINM cannot occur.

**Figure 3.**
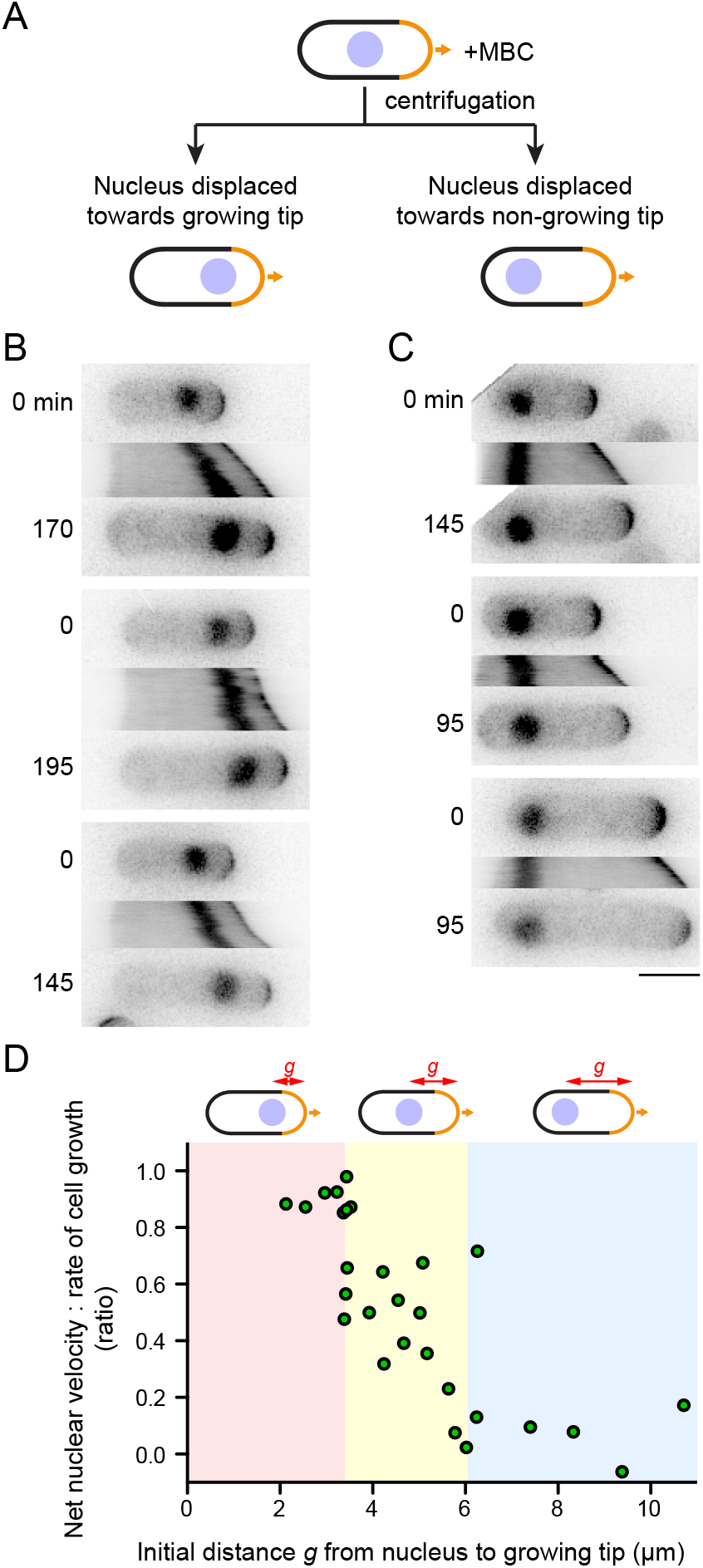
Velocity of MINM depends on distance of nucleus to growing cell tip. **(A)** Schematic of experiment to displace nucleus from cell center by MBC treatment and centrifugation, prior to imaging in presence of MBC. **(B**,**C)** Timepoints and kymographs of nuclear movement from movies of MBC-treated cells in which nucleus is displaced towards growing tip (B) or towards non-growing tip (C). Timepoints (min) correspond to beginning and end of kymographs. Three examples are shown for each condition. **(D)** Net nuclear velocity towards growing cell tips, normalized to rate of cell growth, in cells from these experiments, plotted against distance *g* from nucleus to growing cell tip at start of imaging. Colored regions of graph highlight different types of behavior. Bar for B,C, 5 µm.

### MINM does not require class V myosins

The importance of actin cables in MINM led us to ask whether myosin V motors provide force generation for MINM (see Introduction). Fission yeast contains five myosins: class I myosin Myo1; class II myosins Myo2 and Myp2; and class V myosins Myo51 and Myo52 (East and Mulvihill, 2011). Because Myo1 is associated with cortical actin patches, and Myo2 and Myp2 are associated with the contractile actomyosin ring during cytokinesis, they are unlikely to be involved in MINM and were not considered further. By contrast, class V myosins have broadly conserved roles in intracellular transport (Hammer and Sellers, 2011; Titus, 2018), and therefore we focused on Myo51 and Myo52 (Motegi et al., 2001; Win et al., 2001). Previous work has suggested that Myo51 and Myo52 have distinct functions *in vivo*. In vegetative cells, Myo51 makes minor contributions to interphase actin cable organization and assembly of the actomyosin ring (Huang et al., 2012; Lo Presti et al., 2012; Wang et al., 2014). Myo51 can also move actin filaments relative to each other *in vitro* (Tang et al., 2016), and, although it has no known natural cargos, its motor domain can move an artificial cargo *in vivo* (Wang et al., 2014). Myo52 has a major role in actin-based transport of membrane-associated cargos (Bendezu et al., 2012; Lo Presti and Martin, 2011; Mulvihill et al., 2006; Snaith et al., 2011), and *myo52Δ* deletion mutants have a stubby/spheroid appearance (Motegi et al., 2001; Win et al., 2001). Myo52 is also important for overall actin cable organization, as *myo52Δ* cells display curled and bundled actin cables (Lo Presti et al., 2012). We analyzed MINM in monopolar-growing *myo51Δ* and *myo52Δ* single-deletion mutants and in a *myo51Δ myo52Δ* double mutant (here referred to as *myoVΔ*; (Lo Presti et al., 2012)). In DMSO-treated monopolar-growing *myo51Δ* and *myo52Δ* cells, nuclear movement was similar to wild-type cells, and in MBC-treated *myo51Δ* cells, MINM occurred as in wild-type cells (**Fig. 4A-C;** like *for3* mutants, untreated and DMSO-treated *myo52Δ* and *myoVΔ* cells showed premature bipolar growth, and although enough monopolar-growing cells could be identified in DMSO-treated *myo52Δ* cells to serve as controls, this was not the case for *myoVΔ* cells; see Materials and Methods). Interestingly, in MBC-treated *myo52Δ* cells, nuclear movement was more complex (**Fig. 4D; Movie 5**). In approximately half of the cells (14 out of 29 cells), MINM occurred as in wild-type cells (**Fig. 4D(i)**), indicating that Myo52 is not required for MINM. However, in the remaining cells, (15 out of 29 cells) the nucleus showed no movement for long periods and/or sudden random movements, including movement away from the growing tip (**Fig. 4D(ii)**). In MBC-treated *myoVΔ* cells, MINM occurred essentially as in wild-type cells in 100% of cells (35 out of 35 cells; **Fig. 4F**,**G**; **Movie 6**).

**Figure 4.**
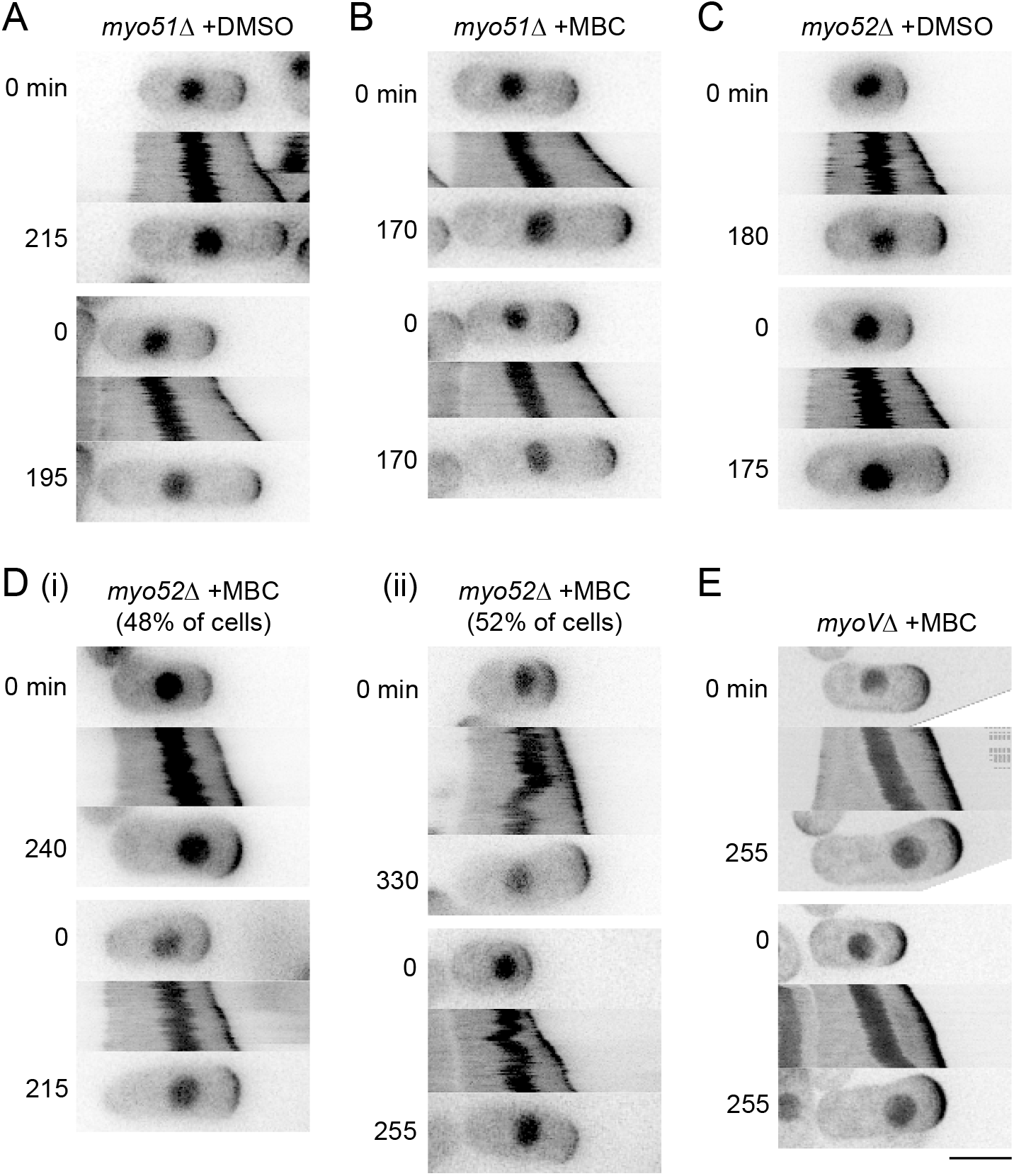
MINM does not require class V myosins Myo51 and Myo52. **(A-E)** Timepoints and kymographs of nuclear movement from movies of cells with the indicated genotypes and treatments. D(i) shows examples of normal nuclear movement in MBC-treated *myo52Δ* cells (48% of cells), while D(ii) shows examples of aberrant nuclear movement, including reversal of direction, in the same population (52% of cells). In E, “*myoVΔ*” indicates double-mutant *myo51Δ myo52Δ*. Because essentially all DMSO-treated *myoVΔ* cells showed premature bipolar growth (see Materials and Methods), no DMSO-treated *myoVΔ* cells are shown. Timepoints (min) correspond to beginning and end of kymographs. Two examples are shown for each condition. Bar for A-E, 5 µm.

These results indicate that neither Myo51 nor Myo52 serves as a motor for MINM. Because MINM depends on actin cables, and actin cables are often disorganized in *myo52Δ* mutants (Lo Presti et al., 2012), we suspect that the aberrant nuclear movement seen in MBC-treated *myo52Δ* cells is due to transient changes in actin cable organization. It is not completely clear why the more complex nuclear movements seen in MBC-treated *myo52Δ* cells were not also observed in *myoVΔ* cells. Because *myoVΔ* cells may have slightly different actin cable organization compared to *myo51Δ or myo52Δ* single mutants (Lo Presti et al., 2012), we speculate that subtle differences in actin organization may be responsible for the observed differences in nuclear movement.

### MINM requires VAPs Scs2 and Scs22

While the actin cytoskeleton represents one possible way to physically link the nucleus with the growing cell tip, another way to provide such a link could be via an endomembrane network involving the NE and the endoplasmic reticulum (ER). The NE is continuous with the ER, and in yeasts, much of the ER is cortical (cortER), closely apposed to the plasma membrane (PM), as a result of ER-PM membrane contact sites that tether cortER to the PM (Manford et al., 2012; Saheki and De Camilli, 2017; Scorrano et al., 2019; Stefan, 2018; West et al., 2011; Wu et al., 2018; Zhang et al., 2012). In both budding and fission yeast, cortER can cover up to 40% of the cytoplasmic face of the PM (Pichler et al., 2001; Pidoux and Armstrong, 1993; Schuck et al., 2009; Zhang et al., 2012). By contrast, in mammalian cells, which have a cortical actin cytoskeleton, less than 5% of the PM is covered by ER-PM contacts.

To investigate whether the NE-ER network contributes to MINM, we analyzed cells lacking the vesicle-associated membrane protein (VAMP)-associated proteins (VAPs) Scs2 and Scs22 (Manford et al., 2012; Murphy and Levine, 2016; Zhang et al., 2012). VAPs are conserved, ER-localized transmembrane proteins involved in lipid transfer, metabolism, signaling, and membrane tethering at membrane contact sites, through interaction with a variety of protein partners on opposing membranes (Gatta and Levine, 2017; Murphy and Levine, 2016; Scorrano et al., 2019; Stefan, 2018; Stefan, 2020). In fission yeast, Scs2 and Scs22 are particularly important for ER-PM contacts, as in *scs2Δ scs22Δ* double-deletion mutants (here referred to as *scsΔ*), the ER shows large-scale detachment from the PM throughout the cell (Ng et al., 2018; Zhang et al., 2012).

In DMSO-treated monopolar-growing *scsΔ* cells, the nucleus moved towards the growing cell tip in 100% of cells (80 out of 80 cells). However, after MBC treatment, there was essentially no nuclear movement in 92% of cells (59 out of 64 cells) (**Fig. 5B**,**C**; **Movie 7**); the remaining 8% of cells showed slight movement, but this did not resemble MINM. These results indicate that, like actin cables, fission yeast VAPs play a critical role in MINM.

**Figure 5.**
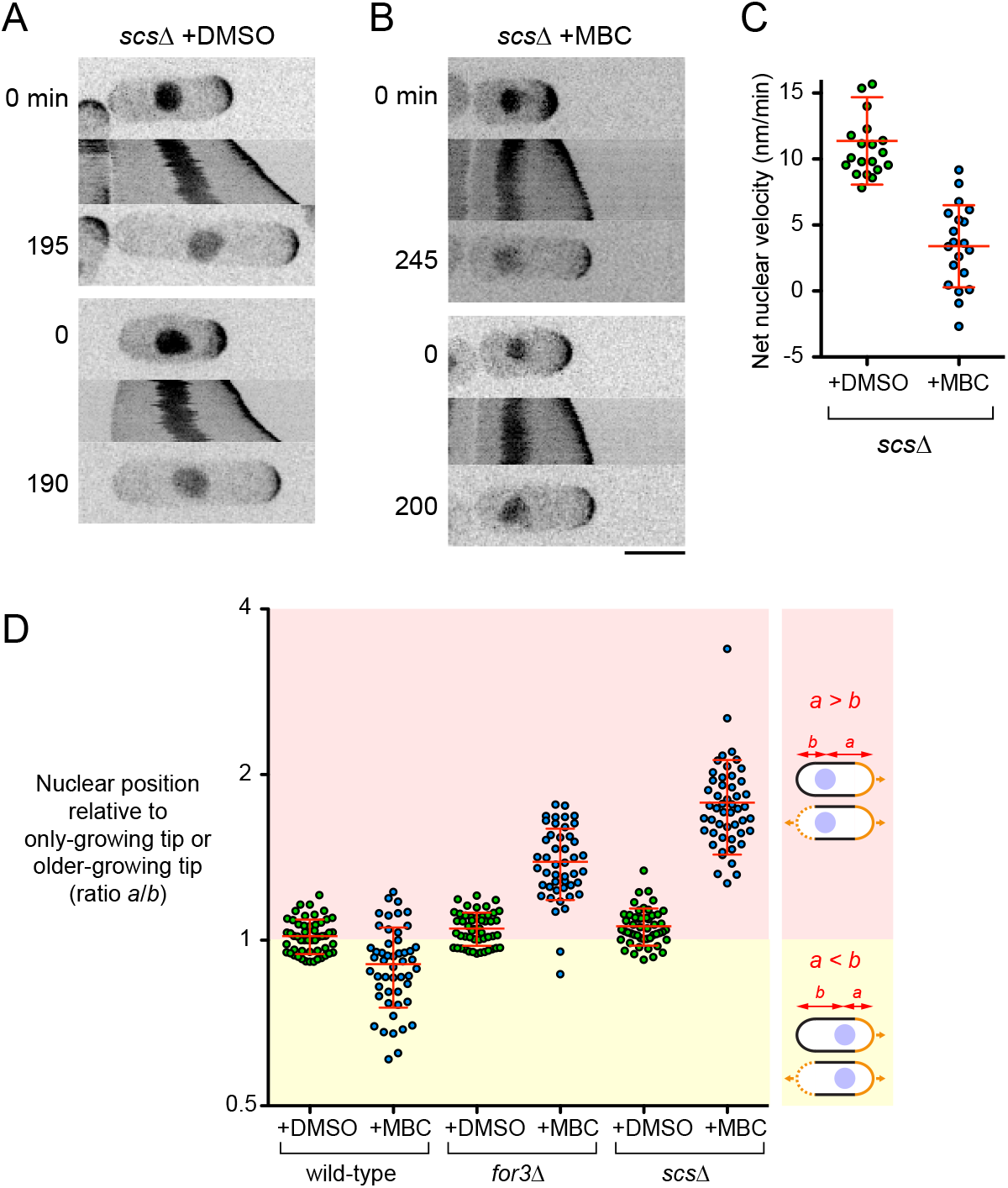
MINM requires VAPs Scs2 and Scs22. **(A, B)** Timepoints and kymographs of nuclear movement from movies of DMSO-treated (A) and MBC-treated (B) double-mutant s*cs2Δ scs22Δ* (“*scsΔ*”) cells. Timepoints (min) correspond to beginning and end of kymographs. Two examples of each condition are shown. **(C)** Net nuclear velocity towards growing cell tips in monopolar-growing DMSO-and MBC-treated *scsΔ* cells. Non-zero mean velocities for *scsΔ*+MBC are likely an artifact of gradual swelling at cell non-growing tips, combined with the method used to determine net nuclear velocity (see Materials and Methods). **(D)** Nuclear position in late-interphase cells with indicated treatments and genotypes. Nuclear position was measured as ratio *a*/*b*, with *a* being the distance from the nucleus to the only-growing cell tip (for monopolar-growing cells; top cell in each diagram) or from the nucleus to the older-growing tip (for bipolar-growing cells; bottom cell in each diagram), and *b* being the distance from nucleus to the other cell tip (see Materials and Methods). For MBC treatments, only monopolar-growing cells were measured, because all cells were monopolar-growing. For DMSO treatments, a mixture of monopolar-and bipolar-growing cells were measured for wild-type and *scsΔ*, while essentially only bipolar-growing cells were measured for *for3Δ*, because *for3Δ* cells show premature bipolar growth (see Materials and Methods).In diagrams, solid orange lines indicate only-growing tips in monopolar-growing cells and older-growing tips in bipolar-growing cells; dashed orange lines indicate newer-growing tips in bipolar-growing cells. Yellow zone corresponds to cells with nuclei closer to only-growing or older-growing tips, while pink zone corresponds to cells with nuclei closer to non-growing or newer-growing tips. Note log2 scale on Y-axis. Red lines show mean and SD. For DMSO-treated cells, differences between wild-type and f*or3Δ* and between wild-type and s*csΔ* were statistically significant (t*-*test, p*=*0.024 and 0.006, respectively). Bar for A,B, 5 µm.

Consistent with both actin cables and VAPs being required for MINM, we found that in MBC-treated *for3Δ* and *scsΔ* mutants, nuclear position in late interphase (i.e. just prior to mitosis) was strongly biased toward the non-growing cell tip (**Fig. 5D**; i.e. in monopolar-growing cells; see Materials and Methods). By contrast, in MBC-treated wild-type cells, in which MINM can occur, nuclear position was biased towards the growing cell tip (**Fig. 5D**).

### Actin cables can buffer MT-based pushing forces on the nucleus

We next asked whether the mechanisms that drive MINM in MBC-treated cells also influence the behavior of the nucleus in unperturbed cells (i.e. in the presence of MTs). In control experiments measuring nuclear position, we noticed that compared to DMSO-treated wild-type cells, the nucleus in DMSO-treated *for3Δ* and *scsΔ* cells showed a small but statistically significant bias towards the non-growing cell tip in monopolar-growing cells and/or the newer-growing cell tip in bipolar-growing cells (**Fig. 5D;** see Materials and Methods). This suggested that MINM-like mechanisms may also be present in untreated cells. If so, then *for3Δ* and *scsΔ* mutants might be expected to show an increased frequency of misplaced septa during normal cell division, because nuclear position in late interphase/early mitosis determines the subsequent plane of cell division (Daga and Chang, 2005). Consistent with this view, we found that while 78% of septa were positioned within 0.02 cell-lengths of the cell center in untreated wild-type cells, this was the case for only 42% and 44% of septa in untreated *for3Δ* and *scsΔ* cells, respectively (**Fig. 6A**,**B**; (Zhang et al., 2012)).

**Figure 6.**
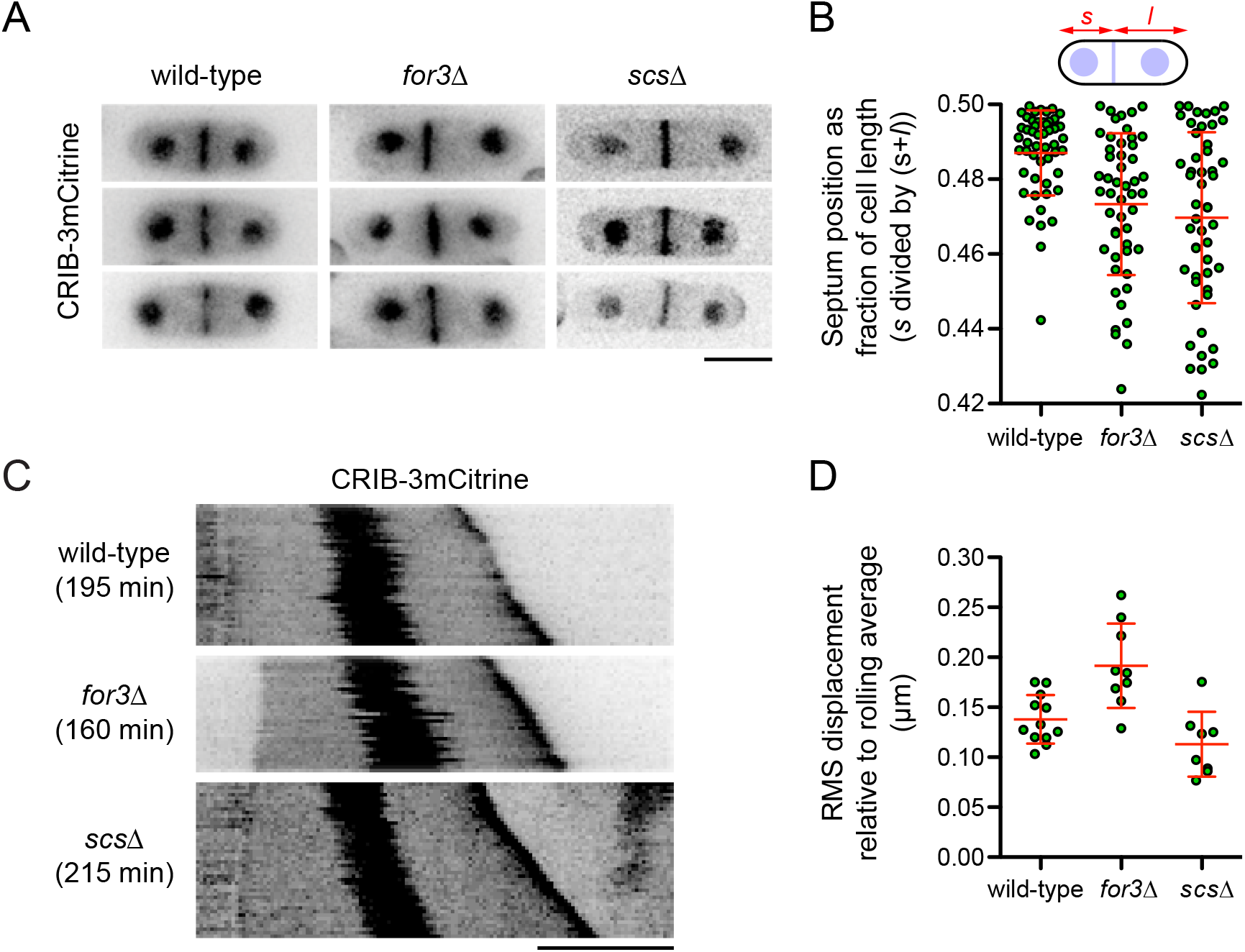
Nucleus and septum positioning defects in *for3Δ* and *scsΔ* mutants. **(A)** CRIB-3xmCitrine images showing septum positioning in untreated cells with the indicated genotypes. Three different cells are shown for each genotype. **(B)** Quantification of septum positioning from images as in A (n=50 cells each). In diagram, *s* and *l* are the shorter and longer distances (respectively) from septum to cell tip. **(C)** Kymo-graphs showing fluctuations in nuclear position in untreated cells with the indicated genotypes. Total time (min) for each kymograph is indicated. **(D)** Root-mean-square (RMS) displacement of nucleus relative to rolling average position over time, from experiments as in C but with higher time resolution (see Materials and Methods and Fig. S2). Each datapoint corresponds to an individual time-lapse movie. Red lines in graphs show mean and SD. Bars, 5 µm.

To complement these experiments, we analyzed fluctuations in nuclear position in untreated interphase cells. Previously, Tran *et al*. showed that MT-based pushing forces lead to fluctuations in nuclear position along the long axis of the cell (Tran et al., 2001), and we observed similar fluctuations in our experiments, (compare for example, **Fig. 1D vs. 1E, or Fig. 2 A vs. 2B**). We hypothesized that if the mechanisms that provide force generation for MINM in MBC-treated cells are also present in untreated cells, then these mechanisms might at least partially resist MT-based pushing forces on the nucleus. We therefore quantified root-mean-square (RMS) displacement of the nucleus from its rolling-average position during growth of untreated wild-type, *for3Δ*, and *scsΔ* cells (see Materials and Methods). Interestingly, and consistent with our hypothesis, RMS displacement of the nucleus in *for3Δ* cells was on average 40-50% greater than in wild-type cells (**Fig. 6C**,**D; Fig. S2)**. To further support these findings, we investigated MT organization in *for3Δ* cells, as it has been suggested that *for3Δ* mutants may have an altered MT distribution (Feierbach and Chang, 2001), and altered MT dynamics could have provided, at least in principle, an alternative explanation for increased fluctuations in nuclear position. However, we found no qualitative or quantitative differences in MT organization between *for3Δ* and wild-type cells (**Fig. S3**). Overall, we interpret these results as indicating that actin cables can normally dampen the transient pushing forces exerted on the nucleus by MTs, such that in the absence of actin cables, fluctuations are increased.

Unlike *for3Δ* mutants, *scsΔ* mutants did not show increased fluctuations in nuclear position compared to wild-type cells (**Fig. 6C**,**D; Fig. S2A)**. One possible reason for this may be that in *scsΔ* mutants, collapse of cortER into cytoplasm **(Zhang et al**., **2012)** could itself dampen MT-based pushing forces on the nucleus (see also Discussion).

## DISCUSSION

Nuclear movement in fission yeast is widely thought to depend on MTs (Dupin and Etienne-Manneville, 2011; Gundersen and Worman, 2013; Xiang, 2018), based on a strong body of evidence that MT-mediated pushing forces are critical for positioning the nucleus in the cell center (Daga et al., 2006; Sawin et al., 2004; Tolic-Norrelykke et al., 2005; Tran et al., 2001). Here we have shown that the fission yeast nucleus can move in the absence of MTs, and we have identified key features of microtubule-independent nuclear movement (MINM). MINM is observed in essentially all cells assayed (i.e. monopolar-growing cells) and is directed towards the growing cell tip. MINM does not exceed the rate of tip growth, and it is strongest when the nucleus is close to the growing tip. MINM requires actin cables but does not require class V myosin motor proteins, which are often employed for actin-based organelle transport (Hammer and Sellers, 2011; Titus, 2018). MINM also depends on the conserved VAP proteins Scs2 and Scs22, which maintain fission yeast cortER via ER-PM contacts (Zhang et al., 2012).

These features suggest that the mechanisms involved in MINM are fundamentally different from previously described mechanisms of nuclear movement (Dupin and Etienne-Manneville, 2011; Gundersen and Worman, 2013; Lele et al., 2018; Xiang, 2018). In nearly all cell types, nuclear movement is driven either by motor proteins acting on MTs or the actin cytoskeleton (“pulling forces”) or by cytoskeletal polymerization itself (“pushing forces”). By definition, MINM is independent of MTs. Our gene-deletion experiments rule out the only plausible actin-based motors, class V myosins, as force generators for MINM. We note that artificial tethering of class V myosin Myo52 to the nucleus has been shown to lead to nuclear translocation in fission yeast (Lo Presti et al., 2012); however, this is non-physiological and occurs at a rate of ∼65 nm/min, about six times faster than MINM. Actin-based pushing forces can also be ruled out as force generators for MINM, because actin cables in fission yeast are nucleated by formins specifically at the growing cell tip, and therefore any actin polymerization-based forces on the nucleus would be expected to oppose rather than support MINM.

Beyond these canonical mechanisms for nuclear movement, it has been shown in mouse oocytes that actomyosin-based vesicle movements generate a pressure gradient that can promote centering of the oocyte nucleus via a mechanism of active diffusion (Almonacid et al., 2015; Brangwynne et al., 2009). However, such a mechanism is unlikely to be involved in MINM, because of the small size of fission yeast relative to mammalian oocytes, as well as fundamental differences in actin organization between the two cell types. Moreover, actomyosin-based vesicle movements in the mouse oocyte depend on class V myosins (Almonacid et al., 2015). Another non-canonical mechanism of nuclear movement involving bulk flow is cytoplasmic streaming, which is generally restricted to very large cells or interconnected cell networks in which advective transport of cellular components may be particularly beneficial (Goldstein and van de Meent, 2015). In animal cell embryos and large plant cells, cytoplasmic streaming is thought to be driven by molecular motors acting on the cytoskeleton (Quinlan, 2016) (Goldstein and van de Meent, 2015; Kimura et al., 2017; Tominaga and Ito, 2015), while in multinucleate filamentous fungi, cytoplasmic streaming has been attributed to hydrostatic pressure gradients, with higher pressure at the center of the colony and lower pressure at hyphal tips (Lew, 2005; Lew, 2011; Xiang, 2018). However, there is no evidence, or apparent benefit, for cytoplasmic streaming in small single-celled fungi, in which small molecules can travel by diffusion. In addition, while cytoplasmic streaming in filamentous fungi is normally closely correlated with rapid hyphal growth (e.g. ∼13-25 µm/min in *Neurospora crassa*; (Lew, 2005; Lew, 2011)), growth rates in fission yeast are much slower (∼0.02-0.04 µm/min at 25°C, depending on specific growth medium; (Baumgartner and Tolic-Norrelykke, 2009; Mitchison and Nurse, 1985)), and cell-tip growth can occur even when MINM cannot (e.g. in *for3Δ* and *scsΔ* mutants). MINM is thus unlikely to be due to cytoplasmic streaming.

While MINM appears to be different from known forms of nuclear movement, the specific mechanisms responsible for MINM force generation remain unclear. Below we discuss two non-exclusive, hypothetical models for this (**Fig. 7**). While both models are consistent with our results, both also contain speculative elements.

**Figure 7.**
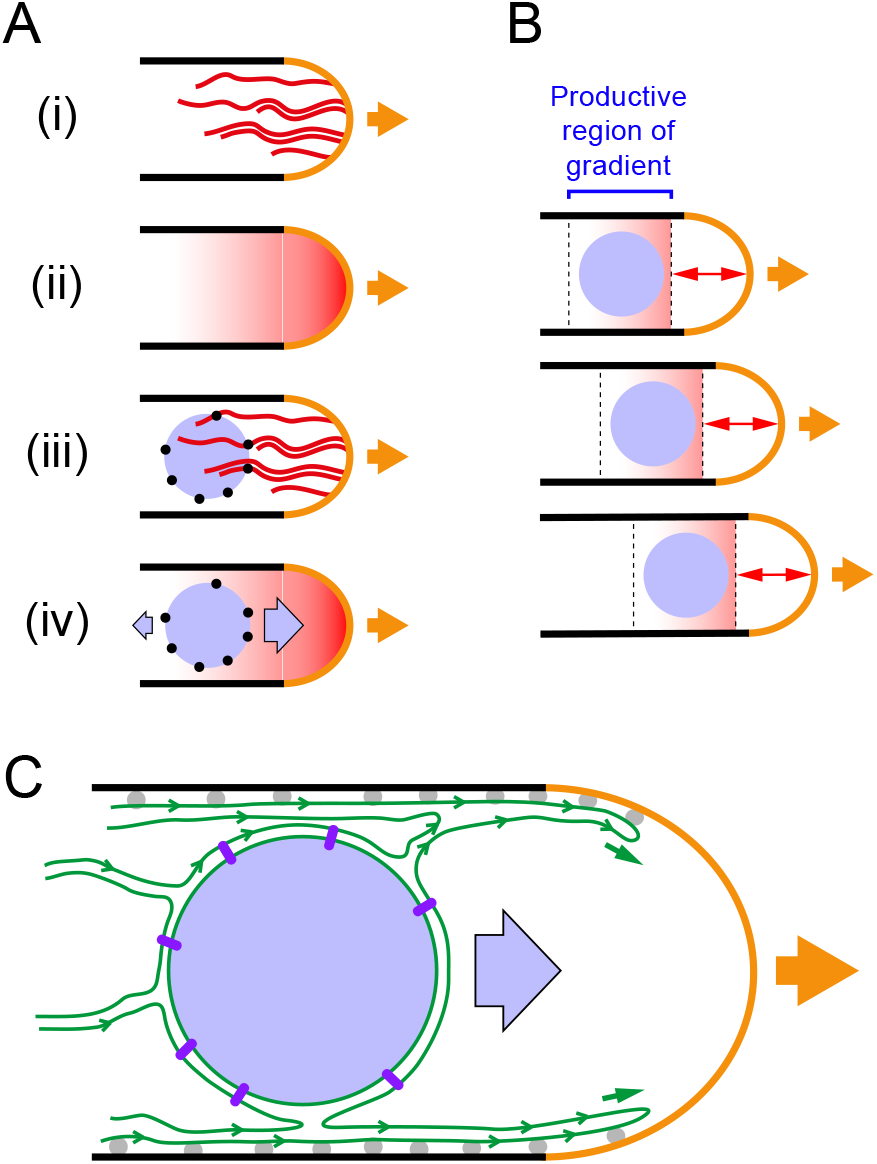
Candidate models for MINM force generation. In all diagrams, cells are drawn growing to the right (orange arrows). **(A)** Model for MINM based on hypothetical nuclear envelope-associated actin filament-binding proteins (NEAF-BPs) and an actin filament-concentration gradient. (i) Cables of dynamic actin filaments (red) are nucleated from the growing cell tip. (ii) Under steady-state conditions, there is an effective time-averaged gradient in the local concentration of actin filaments along long axis of cell. (iii) NEAF-BPs (black circles) on nucleus (light blue) interact with actin filaments. (iv) As a result of NEAF-BPs and the actin filament-concentration gradient, diffusion-driven motion of the nucleus leads to net nuclear movement up the gradient, towards the growing cell tip. See main text for further details. **(B)** Illustration of how MINM could occur even if only a small portion of an actin filament-concentration gradient is productive for diffusion-driven tip-directed movement. Panels show three successive timepoints during cell growth. Because actin filaments are nucleated from the cell tip and turn over as the tip grows, the productive region of the gradient maintains a constant distance from the growing tip (double-headed arrows) and thus moves relative to the deeper cell interior. According to this mechanism, and consistent with observation, the velocity of MINM would not exceed the rate of cell tip growth. **(C)** Model for MINM based on ER membrane dynamics. As the cell grows, ER (green) associated with cell cortex extends forward, as a result of ER-PM contact sites (gray balls). Net flow of ER membrane in direction of cell tip, which could be driven by several factors (see main text), is coupled to MINM via links from NE to nuclear interior (purple rods). Diagrams are schematic and not to scale.

### Model 1

In the first model for MINM force generation, we hypothesize that dynamic, non-motor-based interactions between the nucleus and actin cables could harness the energy of thermal motion to generate directed nuclear movement (**Fig**.**7 A**,**B**). In this scenario, the nucleus would be physically linked to actin cables via NE-associated actin filament-binding proteins (we will refer to such hypothetical proteins as “NEAF-BPs”). Because actin cables are nucleated at the growing cell tip (by formin For3) and contain dynamic filaments of varying length (Martin and Chang, 2006; Wang and Vavylonis, 2008), during steady-state growth (i.e. of monopolar cells) there should be a gradient in the local concentration of actin filaments along the long axis of the cell, with the highest filament concentration close to the growing tip. In the model, random fluctuations in nuclear position would lead to net movement of nucleus in the direction of the growing cell tip, driven by an increased number of interactions of NEAF-BPs with actin filaments. Formally, this type of movement would be similar to haptotaxis of cells in an adhesion gradient (Carter, 1965; Petrie et al., 2009).

One possible objection to this model might be that actin cables may not always be evenly distributed in the region of the cell tip. However, compared to the slow speed of MINM (∼12 nm/min), the flexibility and rapid turnover of actin cables (estimated filament half-life ∼15-20 s (Tang et al., 2014)) may be sufficient to generate a reasonably smooth time-averaged gradient of binding sites. A second potential objection might be that as the nucleus gets very close to the growing cell tip, MINM could be opposed by pushing forces derived from actin polymerization itself; indeed, because actin nucleation occurs at the cell tip, such pushing forces would be greatest close to the tip. However, even if such forces were present, this would not necessarily invalidate the model, because in our experiments the nucleus never actually reaches the cell tip, and mechanism we propose does not require the nucleus to do so. Interestingly, because actin filaments are nucleated from the growing cell tip, the gradient of binding sites for NEAF-BPs would not be stationary with respect to the cell interior but rather would move in the direction of the cell tip at the same rate as cell-tip growth (**Fig. 7B**). Therefore, as long as there is a productive region of the actin filament-concentration gradient within which directed movement of the nucleus can occur (i.e. a “sweet spot”), the model may be plausible.

Currently there are no clear candidates for the NEAF-BPs proposed here. At first glance, likely candidates might include KASH proteins, outer nuclear membrane proteins that interact with inner nuclear membrane SUN proteins to form LINC complexes, which connect the nucleus to the cytoskeleton in many cell types (Burke, 2019; Tapley and Starr, 2013). In metazoan cells, LINC complexes are present over most of the NE. However, in fission yeast, LINC proteins (i.e. KASH proteins Kms1 and Kms2 and SUN protein Sad1) are restricted to the small region of the NE associated with the spindle pole body (i.e. in vegetatively growing cells (Hagan and Yanagida, 1995; Niwa et al., 2000)(Walde and King, 2014)). As this represents only a minute fraction of the nuclear surface, fission yeast LINC proteins are unlikely to fulfil the role suggested for NEAF-BPs. In addition, in preliminary experiments (data not shown) we have found that MINM does not require KASH protein Kms1 (which, unlike Sad1, is nonessential for viability). We have also found (not shown) that MINM does not require inner nuclear membrane protein Ima1 (Hiraoka et al., 2011), the fission yeast homolog of mammalian Samp1, which supports LINC complex interactions with the nuclear interior (Borrego-Pinto et al., 2012).

In a mechanism for MINM involving NEAF-BPs and a gradient of actin filament concentration, the role of ER-PM contacts in MINM could be secondary or indirect. For example, loss of cortER (due to loss of VAPs Scs2 and Scs22) may lead to an increase in cytoplasmic ER (cytoER) sheets/cisternae or tubules, or abnormal cytoplasmic aggregation of ER, which could in turn lead to decreased diffusion of the nucleus, impaired NEAF-BP/actin interactions, or increased resistance against the weak forces driving MINM.

### Model 2

In the second model for MINM force generation, we hypothesize that extension of cortER in the direction of the growing cell tip during cell growth may be a key driver of MINM (**Fig. 7C**). This model is motivated by our finding that Scs2 and Scs22, which are required for ER-PM contacts in fission yeast, are also required for MINM. Because the NE is continuous with cytoER and cortER, ER-PM contacts ultimately help link the NE to the PM. In the model, as the cell tip grows, ER-PM contacts would lead to extension of cortER that would in turn lead to movement of the NE and the nucleus (see below). However, in *scs2Δ scs22Δ* mutants, in which ER-PM connectivity is lost and cortER is collapsed into the cytoplasm (Zhang et al., 2012), cell tip growth would not lead to movement of the NE and nucleus.

In the model, we imagine that during cell-tip growth (i.e. in wild-type cells), extension of cortER could be driven by diffusive spreading of newly synthesized ER membrane (Blom et al., 2011; Jacquemyn et al., 2017) into ER-free regions of the extending PM, followed by “pinning” of new cortER to the PM by VAP-mediated ER-PM contacts (Fig. 7C). In budding yeast, class V myosin Myo4 is important for cortER inheritance from mother cell to bud (Estrada et al., 2003)). However, in fission yeast—perhaps because of differences in cell geometry—class V myosins are not required for cortER extension during cell elongation (although they do contribute to the ability of cortER to reach the extreme cell tip; (Zhang et al., 2012)). This is consistent with our finding that MINM does not require class V myosins.

In order for cortER extension to contribute to MINM, we speculate that as cortER extends and new ER-PM contacts are made, dynamic changes in ER organization may lead to differential membrane tension (i.e. non-homogeneous in-plane tension; (Pontes et al., 2017; Sens and Plastino, 2015)) within the cytoER network. Differential membrane tension should lead to flow of membrane components from lower-tension to higher-tension regions of the network (Dai and Sheetz, 1995). A net tension gradient within the ER network along the yeast-cell long axis (i.e. with higher tension closer to the growing tip) could therefore lead to movement of components of cytoER and of the NE, which itself is linked to the nuclear interior via association of chromatin with nuclear pores and inner nuclear membrane proteins ((Matsuda et al., 2017; Schreiner et al., 2015)). While intracellular membrane tension is more difficult to study than plasma membrane tension, there is evidence for different levels of tension in different intracellular membranes and, more recently, perhaps within the ER itself (Goujon et al., 2019; Pontes et al., 2017; Upadhyaya and Sheetz, 2004). One potential source of differential tension could be the dynamic “tug of war” between sheets and tubules in the ER (Westrate et al., 2015), involving reticulon-, REEP-, and atlastin-family proteins (Wang and Rapoport, 2019); see also (Rangamani et al., 2014). In this context it may be relevant that steady-state maintenance of the ER tubule network requires continuous input of energy, via atlastin-mediated GTP hydrolysis (Powers et al., 2017). Because most lipid synthesis occurs in the ER (Carman and Han, 2011; van Meer et al., 2008), another potential source of differential tension could be nonhomogeneous lipid synthesis within the ER network or, alternatively, dynamic local changes in ER lipid composition (Harayama and Riezman, 2018; Holthuis and Menon, 2014). Regardless of the source, resolution of differential tension via membrane flow might be expected to occur very rapidly (Keren et al., 2008); therefore, for the scenarios proposed here to be plausible, there must be a mechanism to sustain or regenerate differential tension in the longer term (see e.g. (Lieber et al., 2015; Schweitzer et al., 2014)). Given our finding that actin cables are required for MINM, it is possible that motor-independent interactions of ER with actin cables contribute to regulating differential tension and/or its consequences (Prinz et al., 2000). In this context it is particularly interesting that in mammalian cells, immobilization of transmembrane proteins via interaction with the cortical actin cytoskeleton can strongly modulate tension-induced plasma membrane bulk flow (Cohen and Shi, 2020; Shi et al., 2018).

Finally, another possible way for cortER extension to contribute to MINM could be via ER lipid synthesis itself. ER synthesis during cell growth should scale with growth, in order to maintain constant ER density and organization, and if lipids are synthesized essentially homogeneously throughout the ER network during growth, then as the cortER expands unidirectionally in the direction of the growing cell tip, there should be a net flow of cytoER in the same direction. As with membrane flow derived from differential membrane tension, this may be capable of driving MINM. However, in this scenario, it is not immediately obvious how actin cables would contribute to MINM.

### Concluding remarks

Given previous work on nuclear positioning in fission yeast, MINM is both novel and unexpected. Because MINM is slow (∼12 nm/min), it may involve relatively weak forces, and our ability to detect it may depend on the highly polarized geometry of fission yeast growth. It is not yet known whether behaviors analogous to MINM exist in other types of cells. In this context, it is interesting that in the reference frame of the growing cell tip, MINM can potentially be viewed as maintenance of nuclear position relative to that tip. From this perspective, it is possible to imagine that in some cell types, a MINM-like mechanism could contribute to an apparent “fixed” position of the nucleus, as part of a balance of forces. Accordingly, we note that lack of actin cables, which leads to loss of MINM when MTs are absent, leads to increased fluctuations of nuclear position when MTs are present.

Our characterization of MINM also highlights the fact that identifying the source(s) of force generation for intracellular movements is not always straightforward, especially when the associated movements are relatively subtle and/or the forces are unconventional. Recent examples of movement involving unconventional forces include the centering of the mouse oocyte nucleus by active diffusion, mentioned above (Almonacid et al., 2015; Brangwynne et al., 2009), and contraction of cytoskeletal filament networks *in vitro* via entropic forces (Braun et al., 2016; Hilitski et al., 2015; Lansky et al., 2015). How widespread these forces may be in different cell types remains to be determined.

## MATERIALS AND METHODS

### Yeast strain construction and growth

Standard fission yeast techniques were used for strain construction, cell growth, and genetic crosses (Ekwall and Thon, 2017; Petersen and Russell, 2016). Cells were grown as required in YE5S rich medium, using Difco yeast extract (Becton Dickinson), or in Edinburgh minimal medium containing sodium glutamate rather than ammonium chloride as nitrogen source (EMMG, also known as “pombe minimal glutamate”, or PMG; we will refer to this as “minimal medium”). Solid medium used 2% Bacto agar (Becton Dickinson). For antibiotic selection, G418, nourseothricin, and hygromycin were used at 100 µg/mL on YE5S plates. For genetic crosses, cells were mated on SPA plates and incubated for three days at 25°C. Crosses used either random spore analysis or tetrad dissection, and resulting strains were confirmed by colony PCR or fluorescence microscopy, depending on the genotype. A *for3Δ* deletion strain was constructed using PCR-based gene targeting (Bahler et al., 1998) and was confirmed using colony PCR. *Myo52Δ* and *myo51Δ* strains (Win et al., 2001) were obtained from D. Mulvihill, University of Kent, UK. A *myo51Δ myo52Δ* strain (also known as *myoVΔ*) was constructed by deleting myo51 in *a myo52Δ* strain background. *For3-3GFP* and *For3-I930A-3GFP* strains (Martin and Chang, 2006) were obtained from S. Martin, University of Lausanne, Switzerland. A Lifeact-mCherry strain (Huang et al., 2012) was obtained from M. Balasubramanian, University of Warwick, UK. *scs2Δ* and *scs22Δ* strains (Zhang et al., 2012) were obtained from S Oliferenko, Francis Crick Institute, UK. Strains and oligonucleotides used in this study are listed in **Tables S1 and S2**.

### Drug treatments and centrifugal nuclear displacement

To depolymerize microtubules, methyl benzimidazol-2yl carbamate (MBC; Sigma) from a 2.5 mg/mL stock in DMSO was added to cultures at 25°C to a final concentration of 25 µg/mL MBC and 1% (v/v) DMSO (Sawin and Snaith, 2004). For analysis of nuclear movement, conditioned minimal medium containing MBC was added to cells and incubated for 15 mins at 25°C to depolymerize microtubules prior to imaging (in movies, “zero” timepoint represents start of movie, not time of MBC addition). In control experiments, DMSO alone was added to cultures to a final concentration of 1% (v/v).

In preliminary experiments to assess the role of actin cables in MINM, we attempted to use low concentrations of latrunculin A to specifically disrupt actin cables in wild-type cells (i.e. instead of using *for3* mutants; (Lo Presti and Martin, 2011). However, this was found to be unsuitable, as under our long-term imaging conditions, even low concentrations of latrunculin A (e.g. 2 µM) led to impaired cell polarity and cell growth (Mutavchiev et al., 2016).

Nuclear displacement was performed by first depolymerizing microtubules using 25 µg/mL MBC (as above) at 25°C for 15 minutes and then centrifuging cells for three minutes at 13,000 rpm in a microcentrifuge (Carazo-Salas and Nurse, 2006; Daga and Chang, 2005). After centrifugation, all subsequent steps to prepare cells for imaging (see below) were performed in MBC-containing media, to ensure that microtubules remained depolymerized.

### Microscopy

To prepare cells for imaging, freshly-grown cells from YE5S solid medium were used to inoculate a 5 mL minimal medium liquid culture that was incubated overnight at 25°C to mid-log-phase density (0.25-1.0 × 10^7^ cells/mL). Cells were then diluted in 25 mL minimal medium and incubated overnight at 25°C to mid-log-phase density (0.5-1.0 × 10^7^ cells/mL). Approximately 400-600 µL of cell culture, depending on culture density, was then added to a 35mm MatTek coverslip dish (Cat# P35G-0.170-14-C) that had been pretreated by coating with 20 µL of 1 mg/mL solution of soybean lectin (Sigma L1395) in deionized water, air-dried for 10 minutes, and rinsed with deionized water. Cells were allowed to settle on the coverslip dish for 20 min at 25°C. Excess cells were removed using 3-5 washes of 1 mL conditioned minimal medium, which was prepared by growing a large volume of culture (using the same strain as that being imaged) to mid-log-phase density, centrifuging the culture, and recovering the supernatant (Rupes et al., 1999; Sveiczer et al., 1996). After washing, 400 µL of conditioned minimal medium was finally added to the dish, and cells were then imaged. All imaging of cells was performed in conditioned minimal medium.

All imaging was performed at 25°C using temperature-controlled chambers. Nearly all timelapse imaging used a 5 min interval between timepoints; the only exception to this was timelapse imaging of fluctuations in nuclear position, which used a 1 min interval. The majority of live-cell imaging for cells expressing CRIB-3mCitrine (see below for exceptions) was performed on a Deltavision Elite system (Applied Precision) with solid-state illumination, UltimateFocus(tm), and environmental stage control. Imaging was performed using a 100X/1.40 UPLS Apo oil objective (Olympus) and acquired using a Cascade EMCCD camera (Photometrics). For each image, 9 Z-sections were collected (100 ms each), with 0.6 µm spacing. Imaging of mCherry-atb2, GFP-atb2, and Lifeact-mCherry, and of CRIB-3mCitrine in *myo52Δmyo51Δ* (*myoVΔ*) and *scs2Δscs22Δ* (*scsΔ*) double-mutant strains (except for experiments measuring fluctuations in nuclear position in scsΔ) was performed on a customized spinning disc confocal microscope with a Nikon TE2000 microscope base, a modified Yokogawa CSU-10 unit (Visitech) and an iXon+ Du888 EMCCD camera (Andor). The microscope was equipped with a 100x/1.45 NA Plan Apo objective (Nikon), Optospin IV filter wheel (Cairn Research), and MS-2000 automated stage with CRISP autofocus (ASI), controlled using Metamorph software (Molecular Devices). For each image, 11 Z-sections were collected (100 ms each), with 0.6 µm spacing.

Under our imaging conditions, we found that in *for3Δ, for3-I930A-GFP, myo52Δ*, and *myo51Δ myo52Δ* mutants, most control cells (i.e. DMSO-treated cells) displayed premature bipolar growth. While this made it more difficult to identify monopolar-growing cells for analysis of nuclear movement under control conditions, in all cases except *myo51Δ myo52Δ* double mutants, sufficient numbers of cells could nevertheless be identified. To our knowledge, polarity growth patterns in *myo52Δ* and *myo51Δ myo52Δ* mutants have not previously been characterized. With regard to *for3* mutants, the premature bipolar growth we observed is slightly different from the more complex pattern of polarized growth originally described for *for3Δ* cells, in which, after cell division, one daughter cell displays premature bipolar growth, while the other daughter cell shows only monopolar growth (Feierbach and Chang, 2001). While there is no immediately obvious explanation for these different observations, they may be due to differences in preparation of cells before and during imaging, as some live-cell imaging preparations can alter aspects of cell polarity in unintended ways, likely because of mild cell stress (Tay et al., 2018). The imaging protocol used in our work is specifically designed to minimize cell stress (Mutavchiev et al., 2016). In MBC-treated *for3Δ, for3-I930A-GFP, myo52Δ*, and *myo51Δ myo52Δ* cells, monopolar-growing cells were easily identified; we are currently investigating the basis for this different growth pattern.

For measurements of nuclear position in **Fig. 1F**, interphase (non-septating) cells were chosen for measurement from single timepoints in movies, avoiding cells that had immediately finished cytokinesis/septation. Both monopolar-growing and bipolar-growing cells were included in measurements. Distance from the center of nucleus to each of the two cell tips was measured, without regard for whether the tip was growing or not. Graph in **Fig. 1F** shows the ratio *s*/*l*, where *s* corresponds to the shorter of the two distances, and *l* corresponds to the longer of the two distances (i.e. the maximum possible value for *s*/*l* is 1).

For measurements of nuclear position in **Fig. 5D**, cells were chosen for measurement just prior to mitosis/cytokinesis in movies. Depending on the treatment and/or genotype, either monopolar-growing cells, bipolar-growing cells, or a combination of monopolar-growing and bipolar-growing cells were scored. For MBC-treated cells, all cells scored were monopolar-growing. For DMSO-treated wild-type and *scsΔ* cells, a mixture of monopolar-growing and bipolar-growing cells were scored, while for DMSO-treated *for3Δ* cells (most of which showed premature bipolar growth), essentially only bipolar cells were scored. For scoring, septating cells were first identified from movies, and then the movie was stepped back ∼3 frames until a single nucleus was present in the cell (i.e. before karyokinesis). Distance from the center of the nucleus to each of the two cell tips was then measured, taking into account movie data showing which cell tip was the growing tip (for monopolar-growing cells) or had initiated growth first (i.e. “older-growing tip”, for bipolar-growing cells). The distance from nucleus to the growing tip (for monopolar-growing cells) or older-growing tip (for bipolar-growing cells) was termed *a*, and the distance to the other tip was termed *b*. Graph in **Fig. 5D** shows the ratio *a*/*b*. Because this is a ratio, the Y-axis in Fig. 5D uses a log2 scale, and statistical tests were performed on log2-transformed data.

### Image analysis

Image processing and analysis was performed using ImageJ (NIH) and Metamorph (Molecular Devices) software. Movies are shown as maximum projection of Z-sections. Movie timepoints were registered along X- and Y-axes using the “StackReg” plugin of imageJ, using the “rigid body transformation” option. Kymographs were generated by drawing an 11 pixel-wide line across the long axis of the cell and then using the “KymoResliceWide” plugin of ImageJ. All kymographs have a 5 min interval between timepoints.

Nuclear positioning was determined by measuring the distance from the center of the nucleus to both cell tips, using the “Analyze” function of ImageJ. Similarly, septum position was determined by measuring the distance from septum to both cell tips. The shorter distance was termed *s*, and the longer distance *l*, and the ratio *s*/(s+*l*) was plotted. Cell length was calculated using the “Analyze” function of ImageJ. Net nuclear velocity was determined by measuring the distance from the center of the nucleus to the non-growing cell tip at the beginning and at the end of a movie sequence, and then dividing the distance moved by the duration of the movie. Cells were chosen randomly for quantification (for mutants defective in MINM, this meant that cells showing slight movements were also included in quantification). It is important to note that over long time periods, non-growing tips typically show a small amount of swelling and/or local shape changes that do not represent *bona fide* polarized growth. As a result, our measurement method can lead to apparent small but non-zero net nuclear velocities for nuclei that would otherwise be judged (e.g. by eye) to be not moving (see, for example, **Fig. 2B**,**F**,**G**). To avoid any unintended bias in quantification, we did not attempt to correct for these “apparent “non-zero velocities, and the data are simply presented as measured.

Fluctuations in nuclear position were determined from movies using a custom in-house-written ImageJ plugin (DOI: 10.5281/zenodo.3886623). Briefly, time-lapse movies (with 1-minute time interval) are first checked for registration using the ImageJ StackReg plugin, to ensure that the cells to be analyzed are stationary in real space (i.e. apart from their growing cell tips). In the in-house-written plugin, a cut-out image of a user-selected yeast cell is then generated and rotated so that the long axis of the cell is oriented vertically. The cell nucleus is then identified by thresholding, and the centroid coordinates of the nucleus are then measured over time. Values for instantaneous nuclear position in a given cell are then detrended by subtracting a running average of nuclear centroid position in the same cell, so that detrended values fluctuate around zero. The amplitude of fluctuations in a given cell is presented as the root-mean-square displacement of the detrended data.

Numbers of microtubule bundles per cell in wild-type and for3Δ cells were counted manually from still images.

Graphs were plotted using Graphpad Prism (GraphPad, San Diego USA). Movies for presentation were assembled using ImageJ and QuickTime (Apple). For presentation purposes, most movies were rotated, using the “Rotate” function (bicubic) of ImageJ. Figures were made using ImageJ, Photoshop (Adobe) and Illustrator (Adobe).

## ACKNOWLEDGMENTS

We thank Mohan Balasubramanian, Sophie Martin, Dan Mulvihill, Snezhana Oliferenko, and Dan Zhang for strains, and Adriana Dawes, Andrew Goryachev, Alex Mogilner, Snezhana Oliferenko, and members of our laboratory for helpful discussions. This work was supported by the Wellcome Trust ([210659] to KES). SA was supported by a University of Edinburgh Principal’s Career Development PhD Scholarship. The Wellcome Centre for Cell Biology is supported by core funding from the Wellcome Trust [203149].

## DECLARATION OF INTERESTS

No competing interests declared.

---

**Figure S1.**
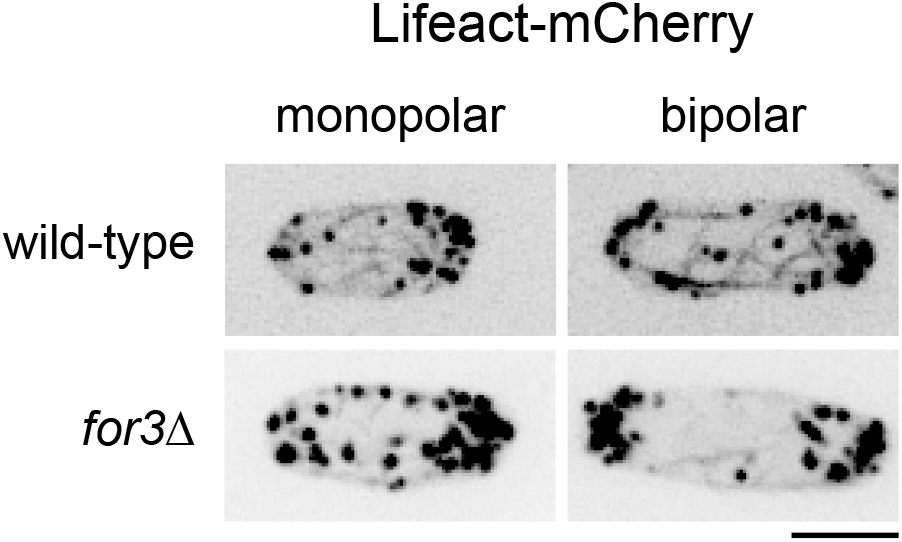
Actin organization in *for3Δ* mutants. Lifeact-mCherry images of interphase monopolar- and bipolar-growing wild-type and *for3Δ* cells. Note that cortical actin patches are much more intense than actin cables. Bar, 5 µm.

**Figure S2.**
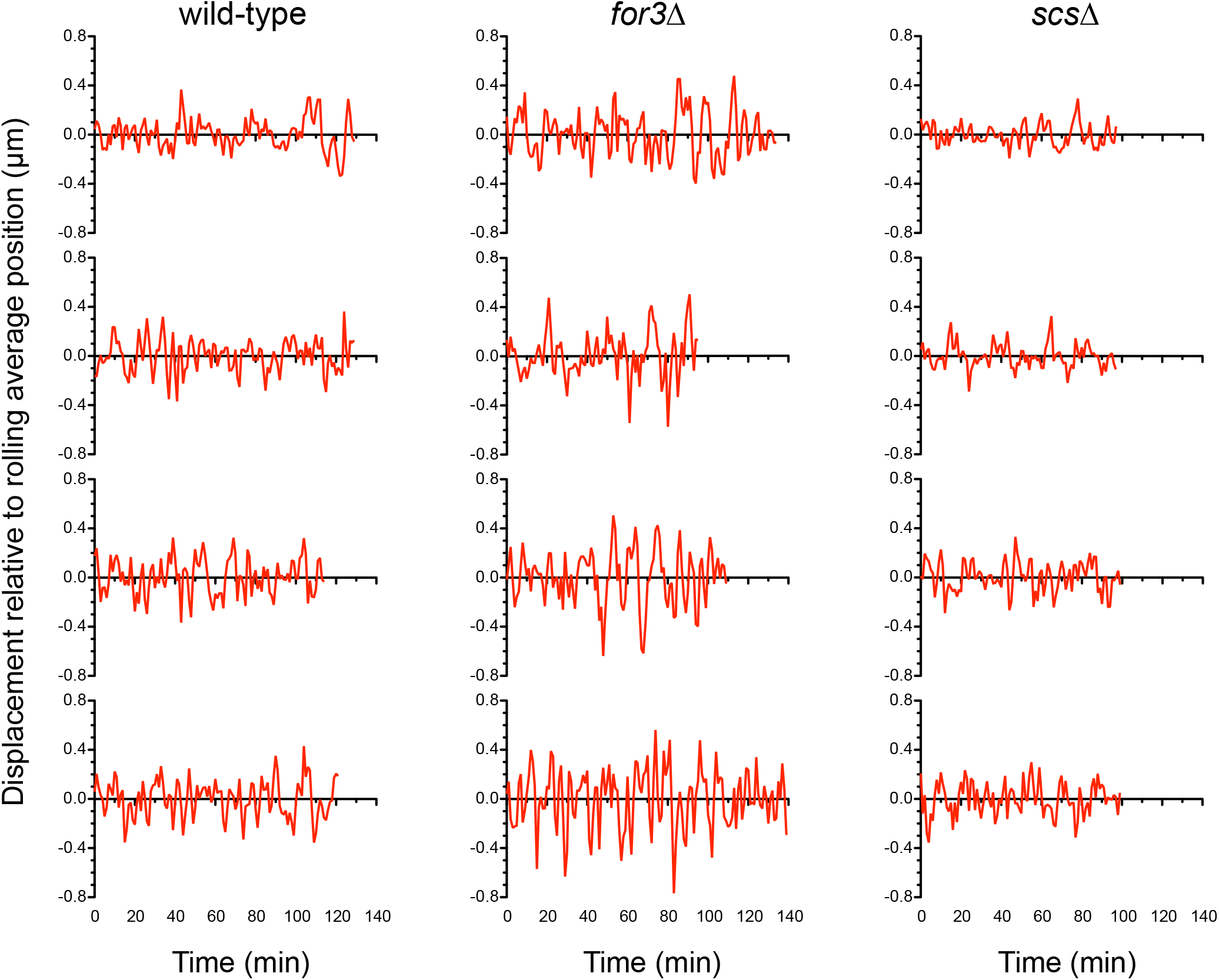
Fluctuations in nuclear position in untreated wild-type, *for3Δ* and *scsΔ* cells. Traces show examples of displacement of the nucleus relative to its rolling average position during normal growth, after detrending (see Materials and Methods). Individual traces correspond to individual root-mean-square (RMS) displacement datapoints in Fig. 6D. Within each genotype, traces are shown from top to bottom in ascending order of RMS displacement; RMS displacements for the examples shown are: 0.125, 0.133, 0.150, and 0.152 µm (wild-type); 0.184, 0.187, 0.221, and 0.240 µm (*for3Δ*); and 0.089, 0.097, 0.123, and 0.132 µm (*scsΔ*). Times of traces range from 96 to 140 min.

**Figure S3.**
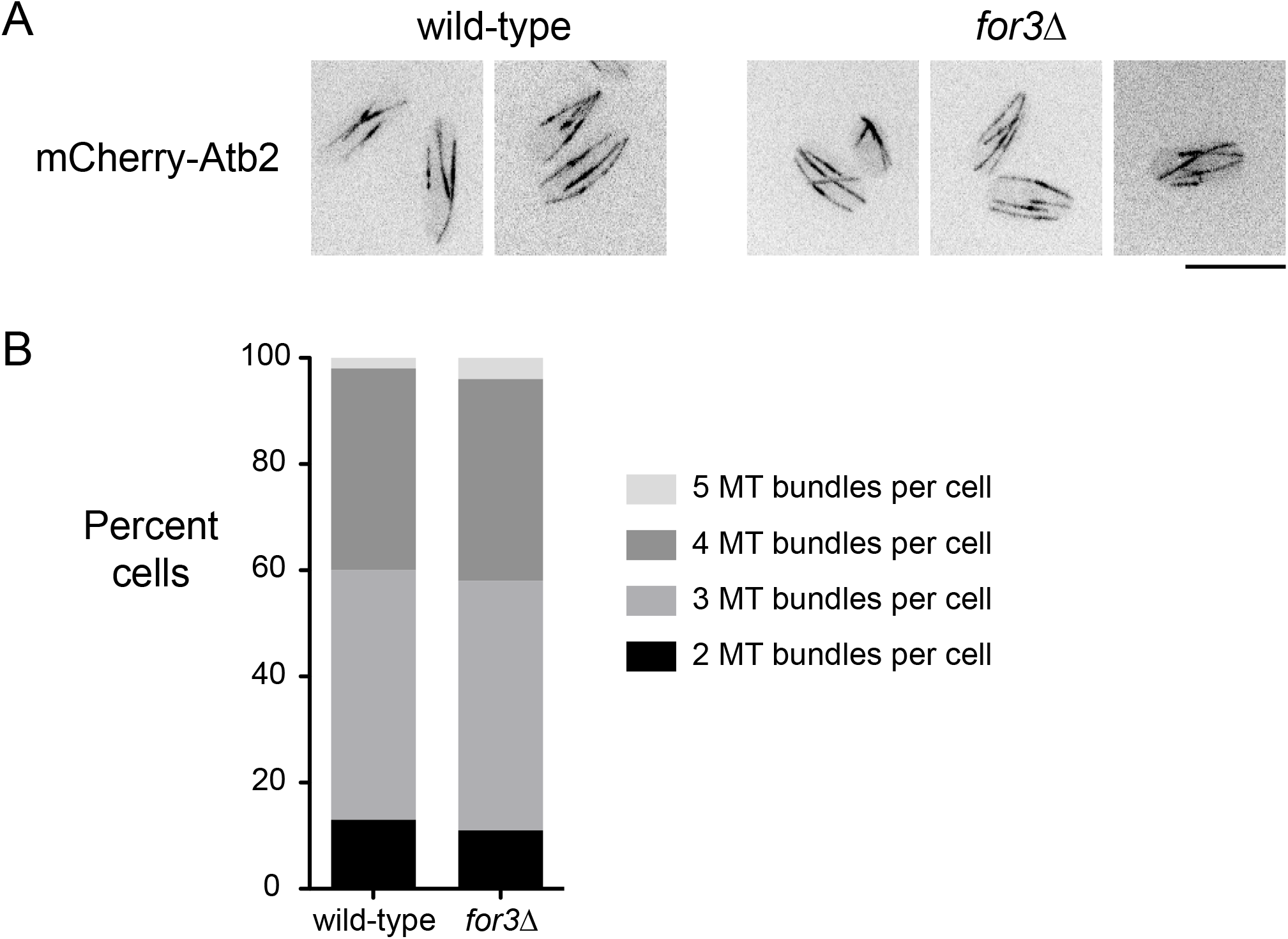
Interphase microtubules in wild-type and *for3Δ* cells. **(A)** mCherry-Atb2 (alpha-tubulin)-labeled microtubules (MTs) in wild-type and *for3Δ* cells. **(B)** Quantification of numbers of microtubule bundles per cell, from images as in A. n=189 wild-type cells and 56 *for3Δ* cells measured. Differences were not statistically significant (chi-square test, p=0.9). Bar, 10 µm.

**Table S1.**
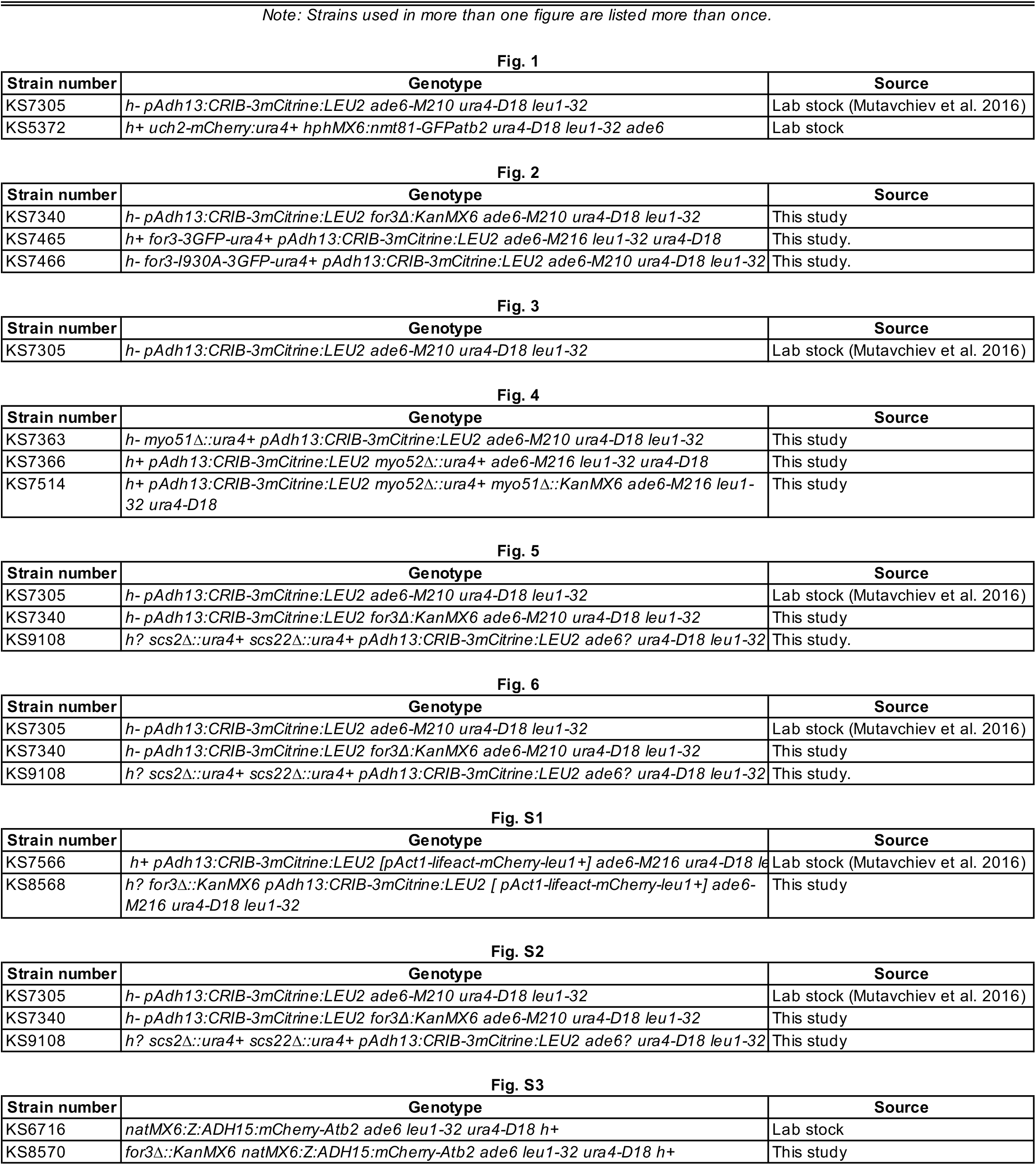
Yeast strains used in this study, listed by figure.

**Table S2.**
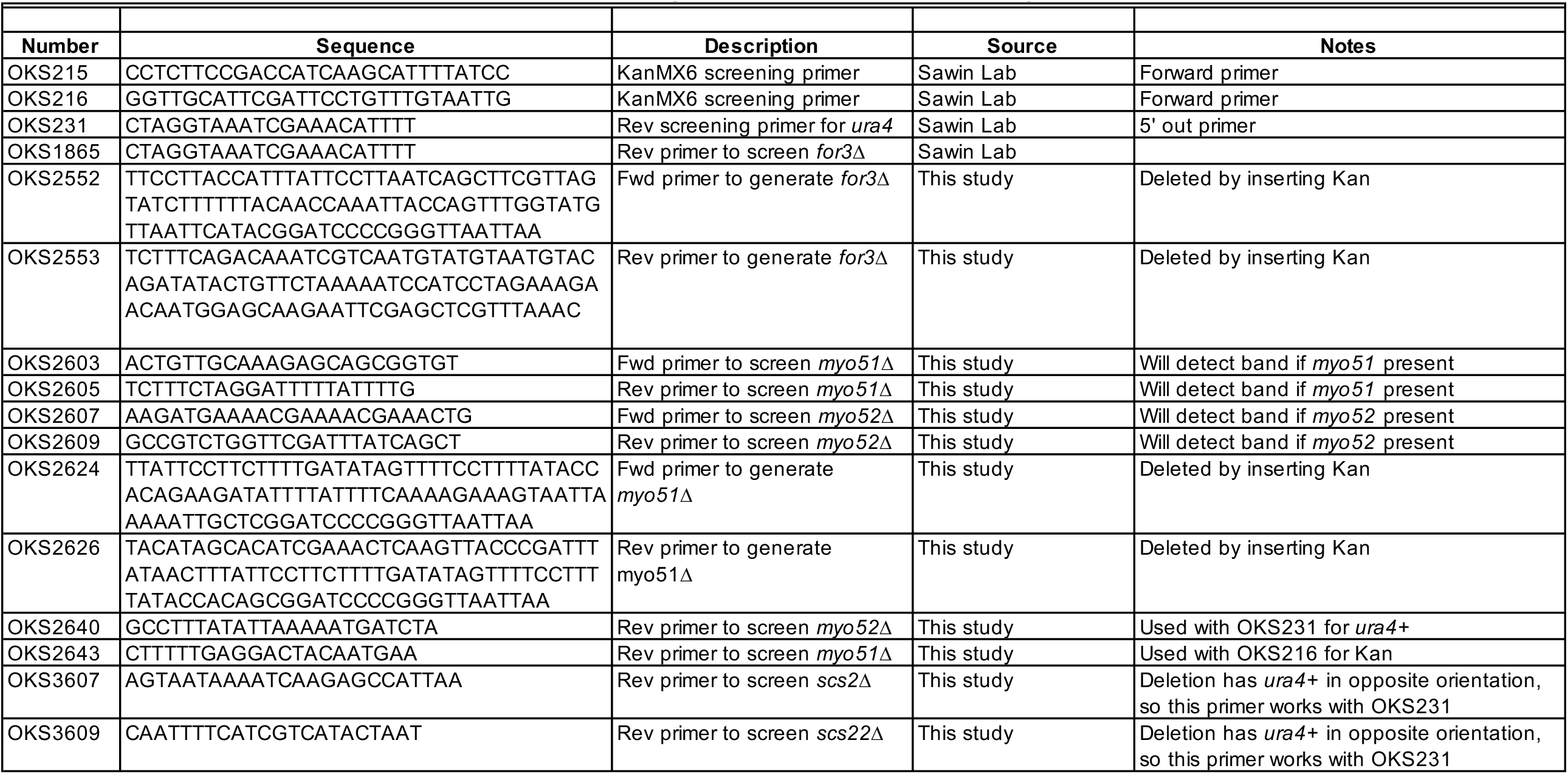
Oligonucleotides used in this study.

## MOVIE LEGENDS

**Movie 1. Microtubules are not required for nuclear movement in fission yeast**. CRIB-3mCitrine in control (+DMSO) and MBC-treated wild-type cells. Cells correspond to those in **Fig. 1D**,**E**. Sequences vary in length. Time interval: 5 min. Total elapsed time of longest sequence: 230 min. Time compression at 15 frames per second playback: 4500x.

**Movie 2. Microtubule-independent nuclear movement (MINM) requires formin For3**. CRIB-3mCitrine in control (+DMSO) and MBC-treated *for3Δ* cells. Cells correspond to those in **Fig. 2A**,**B**. Sequences vary in length. Time interval:5 min. Total elapsed time of longest sequence: 210 min. Time compression at 15 frames per second playback: 4500x.

**Movie 3. MINM requires For3’s actin-nucleating activity**.

CRIB-3mCitrine in control (+DMSO) and MBC-treated *for3-3GFP and for3-I930A-3GFP* cells. Cells correspond to those in **Fig. 2C-F**. Sequences vary in length. Time interval: 5 min. Total elapsed time of longest sequence: 225 min. Time compression at 15 frames per second playback: 4500x.

**Movie 4. Velocity of MINM depends on distance of nucleus to growing cell tip**.

CRIB-3mCitrine in wild-type cells treated with MBC, centrifuged to displace the nucleus, and then imaged in the continued presence of MBC. Cells correspond to those in top panels of **Fig. 3B**,**C**. Sequences vary in length. Time interval: 5 min. Total elapsed time of longest sequence: 170 min. Time compression at 15 frames per second playback: 4500x.

**Movie 5. MINM persists in *myo52Δ* cells, although many cells show additional aberrant nuclear movements**.

CRIB-3mCitrine in control (+DMSO) and MBC-treated *myo52Δ* cells. Middle panels show MINM in *myo52Δ* +MBC. Right-hand panels show aberrant nuclear movement in *myo52Δ* +MBC. Cells correspond to those in **Fig. 4C**,**D**. Sequences vary in length. Time interval: 5 min. Total elapsed time of longest sequence: 330 min. Time compression at 15 frames per second playback: 4500x.

**Movie 6. MINM persists in *myoVΔ* cells**.

CRIB-3mCitrine in MBC-treated *myoVΔ* (i.e. double-mutant *myo51Δ myo52Δ*) cells. Cells correspond to those in **Fig. 4E**. Time interval: 5 min. Total elapsed time: 255 min. Time compression at 15 frames per second playback: 4500x.

**Movie 7. MINM requires VAPs Scs2 and Scs22**.

CRIB-3mCitrine in control (+DMSO) and MBC-treated *scsΔ* (i.e. double-mutant *scs2Δ scs22Δ*) cells. Cells correspond to those in **Fig. 5A**,**B**. Sequences vary in length. Time interval: 5 min. Total elapsed time of longest sequence: 245 min. Time compression at 15 frames per second playback: 4500x.

## Notes

### Competing Interest Statement

The authors have declared no competing interest.

